# Incomplete cerebellar circuit restoration limits functional recovery following SMN therapy in severe spinal muscular atrophy

**DOI:** 10.64898/2026.08.14.744836

**Authors:** Sayan Ruwald, Adela Vankova, Frieda Hanschmann, Christian Menedo, Sandra Wittig, Marie L. Stephan, Vanessa Dreilich, Severine Ruetze, Amy K. Smith, Leonie Sowoidnich, Christian Geis, Stefan Hallermann, Charlotte J. Sumner, Livio Pellizzoni, Beatriz Blanco-Redondo, Florian Gerstner, Christian M. Simon

## Abstract

Spinal muscular atrophy (SMA) is caused by a deficiency in the survival motor neuron (SMN) protein, resulting in degeneration of spinal motor neurons (MNs). However, persistent neurological deficits despite postnatal SMN-restoring therapies suggest that recovery of sensorimotor and supraspinal circuits may be incomplete. The cerebellum has recently emerged as a supraspinal contributor to motor deficits in the severe SMNΔ7 mouse model, yet it remains unclear whether cerebellar pathology is a conserved and therapeutically reversible feature across severe SMA mouse models and clinical subtypes.

Here, we identify cerebellar pathology in Taiwanese SMA mice, characterized by hypoplasia, disrupted organization and loss of Purkinje cells (PCs), altered synaptic circuitry, and impaired cerebellar cortical output. Unlike the previously described p53-dependent PC degeneration in SMNΔ7 mice, cerebellar pathology in Taiwanese SMA mice was associated with developmental disorganization and external granule layer (EGL)-restricted p53 activation. Human cerebellar tissue mirrored this distinction, with p53 activation found in PCs from SMA Type I and in the EGL from SMA Type 0 individuals, indicating that cerebellar pathology arises through distinct mechanisms across severe forms of SMA.

Importantly, two SMN-restoring strategies produced divergent therapeutic outcomes. In SMNΔ7 mice, AAV9-SMN prevented PC degeneration yet incompletely restored cerebellar circuitry. AAV9-SMN-treated Taiwanese mice developed severe ataxia-like deficits, retained profound cerebellar pathology, and survived to approximately one month of age. In contrast, systemic risdiplam rescued cerebellar pathology, motor behavior, and survival in both models.

Together, these findings identify cerebellar pathology as a conserved yet distinct feature across severe forms of SMA and reveal cell type-specific tropism as a critical determinant of therapeutic outcome. More broadly, these findings suggest that successful recovery requires restoration of distributed supraspinal circuit integrity in addition to rescue of spinal motor pathways.

## Introduction

Spinal muscular atrophy (SMA) is an inherited neurodegenerative disorder caused by deficiency of the survival motor neuron (SMN) protein that is characterized by progressive loss of spinal and bulbar motor neurons (MNs), resulting in muscle weakness and paralysis.^1–3^ However, MN loss does not fully account for the complexity of disease manifestations. Increasing evidence indicates that SMA disrupts additional neural circuits across the central nervous system, including spinal sensorimotor and supraspinal systems.^4–17^ Accordingly, therapeutic outcome may depend on preserving not only MN survival but also affected circuit integrity.

Among supraspinal regions, the cerebellum has emerged as a compelling contributor due to its central role in sensorimotor integration, motor coordination, and higher-order behavioral functions.^18^ Previous studies have reported network alterations, hypoplasia, and reduction of Purkinje cells (PCs) - the sole output neurons of the cerebellar cortex - in SMA patients and the SMNΔ7 mouse model.^13,16,19–26^ In this severe SMA mouse model, we recently demonstrated that cerebellar pathology arises intrinsically and independently of spinal sensorimotor dysfunction, revealing a functional dissociation between spinal and cerebellar systems. Consistent with this, SMN deficiency triggers cell-autonomous activation of the p53 pathway in PCs of the SMNΔ7 model, leading to their degeneration. Importantly, selective restoration of SMN in PCs partially improves neurodevelopmental and behavioral abnormalities, including motor deficits.^16^ Notably, another severe SMA model, the Taiwanese mouse, exhibits a similarly severe phenotype and short lifespan but is characterized predominantly by sensorimotor circuit dysfunction despite relatively preserved motor unit integrity.^4,27^ This raises the possibility that dysfunction of neuronal circuits beyond the spinal cord contributes substantially to disease pathogenesis and that cerebellar pathology represents a broader feature of severe SMA. However, whether the cerebellum is affected in the Taiwanese model remains unknown.

The emergence of SMN-restoring therapies, including antisense oligonucleotides, small-molecule splicing modifiers, and viral gene delivery approaches, has transformed the treatment landscape for SMA. However, despite substantial improvements in survival and motor function, neurological deficits frequently persist, indicating incomplete functional recovery.^2,28–31^ Given the emerging role of cerebellar dysfunction in SMA, it remains unclear whether current SMN-restoring therapies effectively re-establish supraspinal motor systems, including the cerebellum, or whether incomplete targeting of the brain contributes to persistent neurological dysfunction.

Here, we investigated cerebellar pathology and the effects of systemic risdiplam treatment and AAV9-mediated SMN gene delivery in two severe SMA mouse models. We demonstrate that cerebellar pathology is a conserved feature of severe SMA that emerges through distinct pathogenic mechanisms and exhibits differential responsiveness to SMN-restoring therapies. Together, these findings identify the cerebellum as a critical determinant of functional recovery and support a broader disease framework in which successful recovery requires restoration of distributed motor circuit integrity.

## Materials and methods

### Study design

The research objectives were to investigate cerebellar pathology in mouse models for SMA and from autopsy tissue of SMA patients. The mice were randomized to treatment groups, and the investigators who assessed the behavioral, histological and electrophysiological outcomes were blinded to the treatment groups. Both male and female mice were included, and data were combined, as no sex-specific differences were observed or have been previously reported in SMA. Data collection was concluded once the predetermined number of animals was reached, based on previous experiments using similar behavioral, functional and histological readouts in SMA mouse models, which consistently yielded robust and reproducible results. The endpoints for animals were selected by previous experiments and literature references.^4,5,8,32^ All data were included if the experiment was technically sound. Each experiment was replicated at least three times in different animals/autopsy tissues with the exception of human tissue due to its limited availability.

### Animal procedures

Breeding and experiments were performed in the animal facilities of Leipzig University (Leipzig, Germany), Johns Hopkins University (Baltimore, MD, USA), and Columbia University (New York, NY, USA). Animal procedures were conducted in accordance with European (Council Directive 86/609/EEC) and German (Tierschutzgesetz) regulations and were approved by the Landesdirektion Sachsen for studies performed at Leipzig University. Experiments conducted at Johns Hopkins University and Columbia University complied with the National Institutes of Health Guide for the Care and Use of Laboratory Animals and were approved by the respective Institutional Animal Care and Use Committees (IACUCs).

Mice were maintained on a 12 h light/12 h dark cycle with ad libitum access to food and water. The following mouse lines were used: SMNΔ7 mice on an FVB background (JAX stock #005025)^33^ and the Taiwanese SMA mouse model (FVB.Cg-Tg(SMN2)2Hung Smn1tm1Hung/J; JAX stock #005058).^34^ Taiwanese SMA mice were originally obtained on a pure FVB background and subsequently backcrossed for seven generations onto a C57BL/6N background. Breeding was performed to generate litters containing approximately 50% SMA and 50% heterozygous control offspring.^35^

The following primers were used for genotyping. SMNΔ7 mice: forward primer (5′– 3′), GATGATTCTGACATTTGGGATG; reverse primers (5′–3′), TGGCTTATCTGGAGTTTCACAA and GAGTAACAACCCGTCGGATTC (wild-type band: 325 bp; Smn knockout band: 411 bp). Taiwanese mice: forward primers (5′–3′), ATAACACCACCACTCTTACTC and GTAGCCGTGATGCCATTGTCA; reverse primer (5′–3′), AGCCTGAAGAACGAGATCAGC (wild-type band: 1050 bp; Smn knockout band: 950 bp).

Adeno-associated virus serotype 9 (AAV9) vectors encoding GFP or human SMN under the control of the GUSB promoter were described previously.^36^ AAV9 vectors encoding GFP or human SMN under the control of the CBA promoter were produced by Vector Builder. For AAV9-mediated gene delivery, postnatal day (P)0–P1 mice were anesthetized by isoflurane inhalation and received a unilateral intracerebroventricular (ICV) injection into the right lateral ventricle containing ∼1 × 10¹¹ genome copies of AAV9 vectors diluted in phosphate-buffered saline (PBS) supplemented with the vital dye Fast Green (Sigma-Aldrich), as previously described.^10,37,38^ SMN-C8 (Roche) and risdiplam (MedChemExpress, HY-109101) were dissolved in sterile dimethyl sulfoxide (DMSO) and administered daily by intraperitoneal (IP) injection at 3 mg/kg.

For phenotypic analysis, treated SMNΔ7 and Taiwanese mice were monitored daily for righting time and body weight, as previously described.^8^ From P20 to P30, body weight and rotarod performance were assessed every 5 days. Each measurement was performed three times and averaged. Righting time was defined as the time required for a pup placed on its back to return to an upright position on all four limbs and maintain this posture for at least 3 s. Tests were terminated after 30 or 60 s, as indicated in the corresponding figure legends. Rotarod performance was defined as the latency to fall from the rotating rod using an accelerating protocol with a cutoff time of 300 s.

### Immunostaining of murine and human tissue

For immunostaining of human cerebellar vermis tissue, samples were obtained following parental or patient informed consent and in accordance with all applicable institutional and ethical regulations. Human tissue was collected during autopsy procedures at Johns Hopkins University (JHU; Baltimore, MD, USA) or obtained from the NIH NeuroBioBank at the University of Maryland (Baltimore, MD, USA). None of the SMA patients had received disease-modifying therapies, with the one exception collected at JHU, as indicated in **Supplementary Table 1**. Human tissue was processed as previously described.^16^ Briefly, cerebellar tissue was cryoprotected in 15% sucrose until the tissue sank, transferred to 30% sucrose overnight at 4°C, embedded in Sakura Tissue-Tek O.C.T. Compound, and frozen in 2-methylbutane cooled with liquid nitrogen. Tissue was sectioned on a Leica CM3050 S cryostat into serial sagittal sections (20 µm) at −20°C and stored at −80°C until use. For immunohistochemistry, sections were incubated for 20 min in Polysciences L.A.B. solution for antigen retrieval at room temperature, washed three times in PBS, and blocked for 90 min in 5% normal donkey serum containing 0.3% Triton X-100 in PBS (PBS-T). Sections were then incubated overnight at 4°C with primary antibodies against parvalbumin to label PCs and phosphorylated p53^S15^ (p-p53^S15^) (**Supplementary Table 2**). The following day, sections were washed six times for 10 min in PBS and incubated for 3 h with Alexa Fluor 488-, Cy3-, or Alexa Fluor 647-conjugated donkey secondary antibodies (Jackson ImmunoResearch; 1:1000 dilution in PBS-T). After six additional washes in PBS, sections were coverslipped using glycerol (3:7) mounting medium.

For immunostaining of murine cerebellum, motor cortex and spinal cord tissue, mice were transcardially perfused with PBS followed by 4% paraformaldehyde (PFA). Tissues were post-fixed overnight in 4% PFA at 4°C. The following day, brains and spinal cords were removed and washed in PBS. The cerebellum was isolated from the remainder of the brain, hemispheres were removed, and one half of the vermis was collected after midsagittal sectioning, as previously established.^16^ For motor cortex analysis, coronal brain sections were collected from the rostrocaudal region located between the olfactory bulb and the anterior forceps of the corpus callosum. Lumbar L1 spinal cord segments were dissected as previously described.^39,40^ Tissue samples were embedded in 5% agar and sectioned on a vibratome (Leica VT1000S) as serial sagittal sections of the vermis (70 µm), coronal sections of the brain (50 µm) or serial transverse sections of the spinal cord (75 µm). Sections were blocked for 90 min in 5% normal donkey serum containing 0.3% Triton X-100 in PBS (PBS-T, pH 7.4). For SMN immunostaining, the blocking solution was supplemented with 3% mouse block (Jackson ImmunoResearch, 715-007-003). Sections were then incubated overnight (or for 2 nights in the case of SMN staining) at room temperature with primary antibodies diluted in blocking solution (**Supplementary Table 2**). The following day, sections were washed six times for 10 min in PBS and incubated for 3 h with Alexa Fluor 488-, Cy3-, or Alexa Fluor 647-conjugated donkey secondary antibodies (Jackson ImmunoResearch; 1:1000 dilution in PBS-T). After additional 6 times for 10 min washes in PBS, sections were mounted on glass slides in glycerol:PBS (3:7), as previously described.^8,39^

For immunostaining of neuromuscular junctions (NMJs), mice were transcardially perfused, and muscles were dissected and post-fixed in 4% PFA for 2 h before transfer to PBS. The following day, single muscle fibers were teased apart and washed three times for 10 min in PBS. Postsynaptic acetylcholine receptors were labeled with Alexa Fluor 555-conjugated α-bungarotoxin (BTX) for 20 min. Subsequently, muscle fibers were washed five times for 10 min in PBS and blocked in 5% donkey serum containing 0.3% Triton X-100 in PBS-T for 1 h. Primary antibodies against neurofilament (NF) and synaptic vesicle protein 2 (SV2) were applied overnight at 4°C to visualize presynaptic nerve terminals (**Supplementary Table 2**). The following day, fibers were washed three times for 10 min in PBS and incubated with appropriate secondary antibodies for 1 h at room temperature. Finally, muscle fibers were washed three times for 10 min in PBS and mounted on glass slides in glycerol:PBS (3:7), as previously described.^4,41^

### Confocal imaging and analysis

For PC quantification and lobule-specific analysis, sagittal 70 µm murine or 20 µm human vermis sections were scanned using a 10x or 20x objective. Z-stacks were acquired at 4.0 µm intervals to capture PC somata. Individual image planes were automatically stitched using Leica LAS X software to generate a complete image of each vermis section. For quantification, each stack of murine tissue was divided into two smaller stacks (∼35 µm), and measurements were averaged between stacks. Quantitative analysis was performed using Leica LAS X software. The area of the entire vermis and individual lobules was determined manually. The molecular layer, PC layer, granule cell layer, and white matter were similarly outlined and quantified for each of the ten cerebellar lobules. Three vermis sections were analyzed per animal as previously established.^16^

For MN quantification, 11 transverse sections (75 µm) of the L1 spinal cord segment were scanned using a 20x objective. MNs were counted from z-stack images acquired at 4 µm intervals throughout the entire spinal cord segment. Only ChAT-positive MNs located within the ventral horn and containing a visible nucleus were included to avoid double counting across adjacent sections, as previously described.^4,40,42^

For synaptic density and gemini of coiled bodies (GEM) quantification, images were acquired at z-step intervals of 0.4 µm for synapses and 1 µm for GEMs using a 63× oil-immersion objective. The Leica LAS X z-compensation function was used to adjust laser intensity throughout the z-stack and maintain consistent signal quality. A minimum of two image stacks were acquired per animal. Synaptic density was quantified using Leica LAS X software. For each lobule, at least four PCs, and for each spinal cord section at least ten MNs, were randomly selected for analysis of VGLUT1⁺, VGLUT2⁺, and VGAT⁺ synapses. Synapses were manually quantified on the soma and on dendritic segments located 0–50 µm from the soma, as previously described.^39,42^ To minimize background-related errors, only synaptic puncta visible in at least two consecutive optical sections were included in the analysis.

For motor cortex analysis, coronal brain sections containing the motor cortex were identified using anatomical landmarks and corresponding reference sections from the Allen Mouse Brain Atlas (Paxinos and Franklin’s The Mouse Brain in Stereotaxic Coordinates, eBook ISBN: 9780128161944). Specifically, sections were selected based on the presence of the characteristic cortical indentation dorsal to the piriform cortex and the appearance of the orbital cortex in ventral cortical regions. Images were acquired from the dorsolateral motor cortex. For each image, z-stacks spanning 50 µm were acquired with a step size of 3 µm. Four optical planes were selected from each z-stack for quantification, resulting in four counts per motor cortex image. Images were cropped to a standardized area of 500 × 500 µm before analysis using ImageJ. CTIP2- and SMI-32-positive neurons in layer V were classified as pyramidal neurons and manually quantified in ImageJ. Cell counts were obtained from four optical planes per section and averaged for each animal.

For the analysis of muscle innervation, a minimum of 200 randomly selected NMJs per muscle sample were scanned at 3 μm z-steps and quantified for each biological replicate. Only BTX^+^ endplates that lack pre-synaptic coverage by both SV2 and NF were scored as fully denervated as previously described.^32^

### Whole-cell patch-clamp slice recordings of Purkinje cells

Whole-cell patch-clamp recordings from PCs were performed in P10 Taiwanese mice as previously described.^16^ Following decapitation, the cerebellum was rapidly removed and immersed in ice-cold, oxygenated (95% O₂/5% CO₂) sucrose-based artificial cerebrospinal fluid (aCSF) containing (in mM): 2.5 KCl, 1.1 CaCl₂, 4 MgCl₂, 25 NaHCO₃, 1.25 NaH₂PO₄·H₂O, 10 D-glucose, and 220 sucrose. Parasagittal cerebellar slices (300 µm) were prepared using a vibratome (HM 650 V, Microm, Thermo Fisher Scientific, UK) and transferred to a holding chamber containing recording aCSF composed of (in mM): 125 NaCl, 2.5 KCl, 1.25 NaH₂PO₄·H₂O, 26 NaHCO₃, 3 D-glucose, 1.1 CaCl₂·H₂O, 1 MgSO₄·7H₂O, and 17 sucrose (310 mOsm, pH 7.3, equilibrated with carbogen). Slices were incubated at 35°C for 30 min and subsequently maintained at room temperature (21–23°C) until use.

For recordings, slices were continuously perfused with oxygenated aCSF at room temperature. PCs were identified by their location within the PC layer, large soma, and characteristic dendritic arbor extending into the molecular layer. Patch pipettes (4–7 MΩ) were pulled from borosilicate glass (GB200F-10, Science Products, Hofheim am Taunus, Germany) using a Flaming-Brown puller (P-97, Sutter Instruments, CA, USA). Pipettes were filled with an intracellular solution containing (in mM): 150 K-D-gluconate, 10 NaCl, 10 HEPES, 3 Mg-ATP, 0.3 Na-GTP, and 0.05 EGTA (pH 7.3; 290–300 mOsm/kg H₂O).

Recordings were obtained in current-clamp or voltage-clamp mode using a HEKA EPC10/2 amplifier (HEKA Elektronik, Lambrecht/Pfalz, Germany), digitized at 20 kHz, and acquired with Patchmaster software (HEKA Elektronik). Signals were low-pass filtered at 3 kHz. Resting membrane potential (RMP) and spontaneous activity were recorded during a 10 s baseline period without current injection. Only cells with an RMP ≤ −45 mV, AP amplitudes ≥ 30 mV, and the ability to generate at least 10 APs during sustained depolarization were included in the analysis.^16^ Passive membrane properties were assessed in current-clamp mode by holding cells at −80 mV to suppress spontaneous firing and applying alternating hyperpolarizing and depolarizing current steps (300 ms; 4 pA increments). Frequency–current (F–I) relationships were determined using 1 s depolarizing current injections in 10 pA increments. Data were analyzed offline using Patchmaster software.

### Western blot analysis

Whole murine spinal cord and cerebellum tissue were prepared in 1x LDS buffer (Invitrogen) and resolved using the NuPAGE precast gel system (Invitrogen) by SDS-PAGE. Extracts (20 µg) were run on Novex Bis-Tris 12% gels and transferred onto an iBlot2 transfer stack nitrocellulose membrane (Invitrogen) using the iBlot2 Dry Blotting system unit (Invitrogen). After protein transfer, the membranes were blocked for 1 h in 5% non-fat dry milk prepared in 1 x PBS. The membranes were then incubated with the corresponding antibodies overnight. Thereafter, the membranes were incubated with IRDye 680RD or 800CW secondary antibodies (Li-cor) followed by visualization using a near-infrared imager (Odyssey; Li-cor).^43^ Band intensities were analyzed by Fiji ImageJ.

### Statistics

In this manuscript, N refers to the number of patients or mice in each group, and n refers to the number of cells analyzed. Sample sizes (N and n) for each experiment are indicated in the corresponding figure legends. Results are presented as mean ± standard error of the mean (SEM). Normality was assessed using the Shapiro–Wilk test. For comparisons between two groups, parametric data with equal variances were analyzed using an unpaired Student’s t-test or multiple unpaired t-tests. When variances were unequal, Welch’s t-test was applied. Nonparametric data were analyzed using the Mann–Whitney test or multiple Mann–Whitney tests. For comparisons among three or more groups, parametric data were analyzed using one-way ANOVA followed by Tukey’s multiple-comparison test. Nonparametric data were analyzed using the Kruskal–Wallis test followed by Dunn’s multiple-comparison test. When variances were unequal, Welch’s ANOVA followed by Dunnett’s T3 multiple-comparison test was used. Time-course experiments involving more than two groups were analyzed using two-way ANOVA followed by Tukey’s multiple-comparison test or mixed-effects model (REML). Survival curves were compared using the log-rank (Mantel–Cox) test. The statistical tests used for each experiment are indicated in the corresponding figure legends. All statistical analyses were performed using GraphPad Prism 10 (GraphPad Software), and exact p-values are reported within the figures.

## Results

### Defective cerebellar development in the Taiwanese mouse model and the most severe form of human SMA pathology

Our previous studies established robust cerebellar pathology in the severe SMNΔ7 mouse model that contributes to motor dysfunction, whereas a milder SMA model lacks overt cerebellar abnormalities.^16^ To determine whether this phenotype is model-specific or represents a conserved feature of severe SMA, we examined the Taiwanese SMA mouse model at end stage (P10). Neurofilament immunostaining of sagittal brain sections revealed a pronounced reduction in cerebellar size (**Fig. 1A**, upper panel). To extend our assessment to other major supraspinal component of the motor system, we next evaluated the integrity of the motor cortex. In contrast to the cerebellum, neurofilament and CTIP2 immunostaining^44^ demonstrated preserved motor cortex morphology and unchanged density of layer V corticospinal projection neurons (**Fig. S1A, B**), indicating selective vulnerability of the cerebellum among supraspinal motor structures.

**Figure 1.**
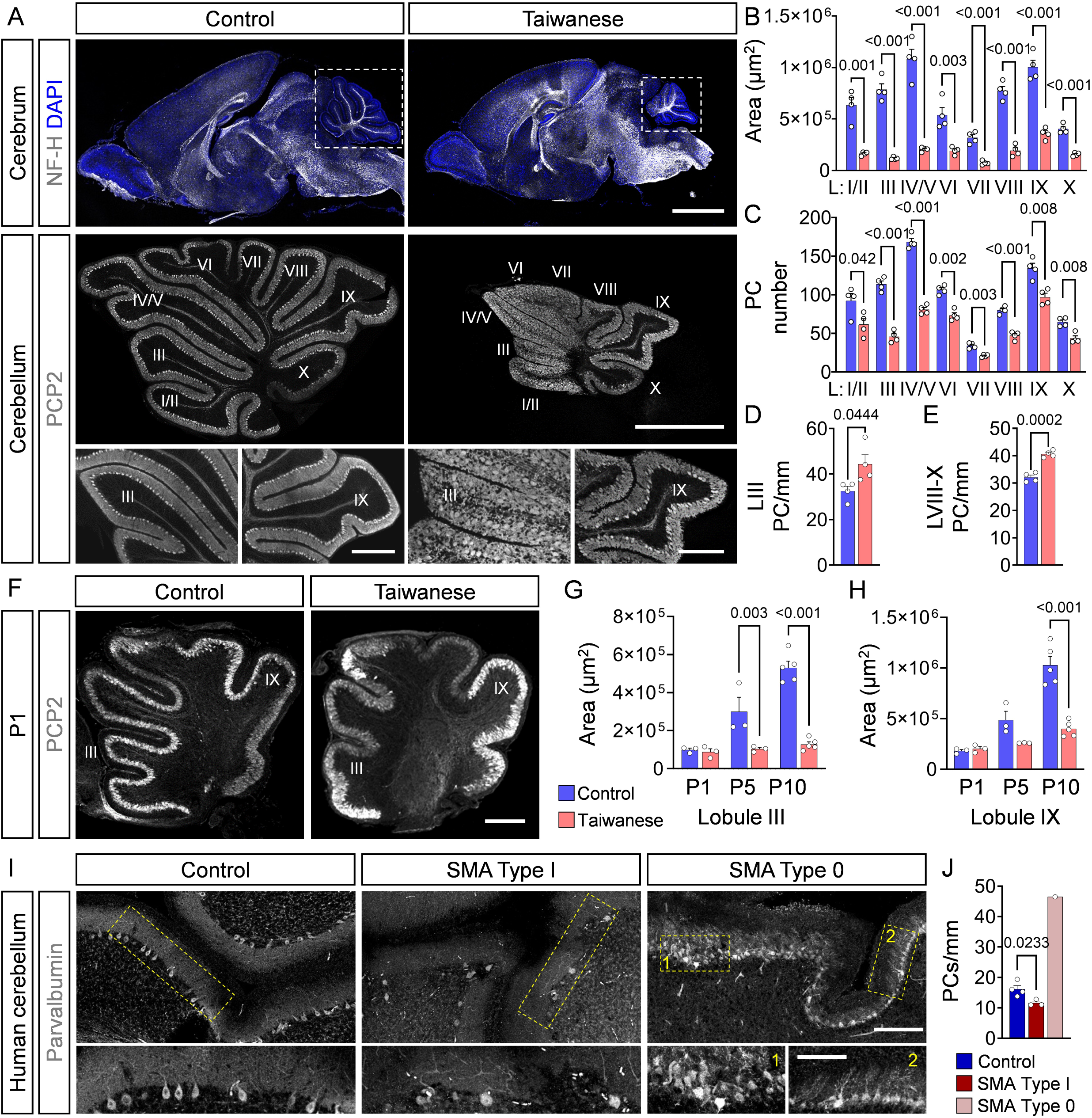
Developmental cerebellar pathology in the Taiwanese SMA mouse model mirrors severe human SMA. **(A)** Top: Confocal images of sagittal whole-brain sections immunostained for neurofilament H (grey) and DAPI (blue) in P10 control and Taiwanese SMA mice. Scale bar, 2 mm. Middle: Confocal images of sagittal cerebellar sections showing PCP2⁺ Purkinje cells (PCs) across lobules (L) I–X. Scale bar, 1 mm. Bottom: Higher-magnification images of lobules III and IX. Scale bars, 500 μm (control) and 200 μm (Taiwanese SMA). **(B)** Lobule-specific quantification of total cross-sectional area. **(C)** Lobule-specific quantification of absolute PC numbers. **(D, E)** Quantification of PC density in lobules III (D) and VIII–X (E). N = 4 control and N = 4 Taiwanese SMA mice for all analyses shown in panels B–E. (**F**) Confocal images of sagittal cerebellar sections showing PCP2⁺ PCs of P1 control and Taiwanese SMA mice. Scale bar, 200 µm. (**G, H**) Quantification of the area of lobules III (G) and IX (H) of P1, P5, and P10 control and Taiwanese SMA mice. N = 3–4 mice per genotype and age. **(I)** Representative confocal images of sagittal human cerebellar sections showing parvalbumin⁺ PCs from a control (0.8 months), SMA Type I (2 months), and SMA Type 0 (0.5 months) patient. Scale bar, 200 μm; inset, 50 μm. **(J)** Quantification of PC density in control (N = 4), SMA Type I (N = 3), and SMA Type 0 (N = 1) cerebella. Statistical analyses were performed using multiple unpaired t-tests (B, C, G, H) and unpaired Student’s t-tests (D, E and control versus SMA Type I in J). The SMA Type 0 sample is shown for descriptive comparison only and was not included in statistical analyses.

To characterize cerebellar morphology in detail, sagittal vermis sections were immunolabeled for Purkinje cell protein 2 (PCP2), a specific marker of PCs,^45^ and analyzed by confocal microscopy (**Fig. 1A**, middle and lower panels). This confirmed a marked reduction in cerebellar area and an approximately 40% decrease in total PC number across all lobules in end stage Taiwanese mice (**Fig. 1A–C** and **Fig. S1C**). Despite the reduction in PC number, PC density across anterior and posterior lobules was increased by approximately 25%, reflecting the marked reduction in cerebellar size (**Fig. 1D, E** and **Fig. S1D**).

In contrast to the homogeneous increase in density, PC organization differed strikingly along the anteroposterior axis. Posterior lobules VIII–X retained a largely preserved cerebellar architecture, with proportionally reduced but well-defined cortical layers and a continuous monolayer of PCs comparable to age-matched control mice (**Fig. 1A** and **Fig. S1E–H**). In contrast, PCs in the anterior and intermediate lobules failed to form a continuous PC layer and instead remained scattered throughout the cerebellar cortex (**Fig. 1A**), resembling the immature organization observed during early postnatal development (**Fig. 1F**).

To determine whether this regional disorganization reflected impaired cerebellar growth, we quantified lobule size throughout postnatal development. Although anterior lobules exhibited less foliation, their size was largely preserved in P1 Taiwanese mice (**Fig 1F-H**). In contrast, postnatal expansion was markedly impaired in Taiwanese mice, resulting in progressive hypoplasia throughout anterior and posterior lobules (**Fig. 1F-H**). Taken together, these findings identify developmental cerebellar hypoplasia as a defining feature of the Taiwanese SMA model.

To determine whether the divergent cerebellar phenotypes observed in severe SMA mice are reflected in human disease, we analyzed cerebellar tissue from one SMA Type 0 patient (0.5 months) and compared it with control tissue (0.8–3.1 months) as well as SMA Type I samples (2–7 months). Consistent with our earlier observations,^16^ a new series of immunostainings confirmed an approximately 30% reduction in parvalbumin-positive (PV⁺) PC number in SMA Type I patients relative to controls (**Fig. 1I, J**), recapitulating the PC degeneration previously described in SMNΔ7 mice.^16,21^ In contrast, although limited to randomly sampled cerebellar regions from a single SMA Type 0 patient due to the scarcity of available tissue, PC density appeared approximately threefold higher than in controls (**Fig. 1I, J**). Because PC density remained stable across the analyzed age range and postmortem intervals in both control and SMA Type I samples (**Fig. S1I**), this observation is unlikely to be explained by these variables. Importantly, focal regions displayed disrupted PC layer organization with misaligned PCs (**Fig. 1I, box 1**), closely resembling the structural abnormalities observed in Taiwanese mice.

Together, these findings establish Taiwanese mice as a severe SMA mouse model characterized by developmental cerebellar hypoplasia and regional PC layer disorganization, key architectural abnormalities that are also evident in the human cerebellum from one SMA Type 0 individual.

### Impaired dendritic architecture, synaptic organization, and PC output in Taiwanese SMA mice

To determine whether the structural abnormalities observed in Taiwanese SMA mice impair cerebellar circuit function, we next analyzed PC synaptic organization and functional properties. First, we performed synaptic analyses in lobule VI to facilitate direct comparison with the previously characterized SMNΔ7 model^16^. Remarkably, the dendritic arbor of Taiwanese PCs was profoundly reduced (**Fig. 2A, C**), precluding reliable quantification of dendritic synapses and restricting analyses to somatic inputs. While VGLUT2⁺ climbing fiber inputs were unchanged **(Fig. S2A, B)**, both VGLUT1⁺ excitatory and VGAT⁺ inhibitory synapses associated with PC somata were increased **(Fig. 2A-D)**, consistent with a redistribution of synaptic contacts toward the soma caused by the markedly reduced dendritic arbor.^46,47^

**Figure 2.**
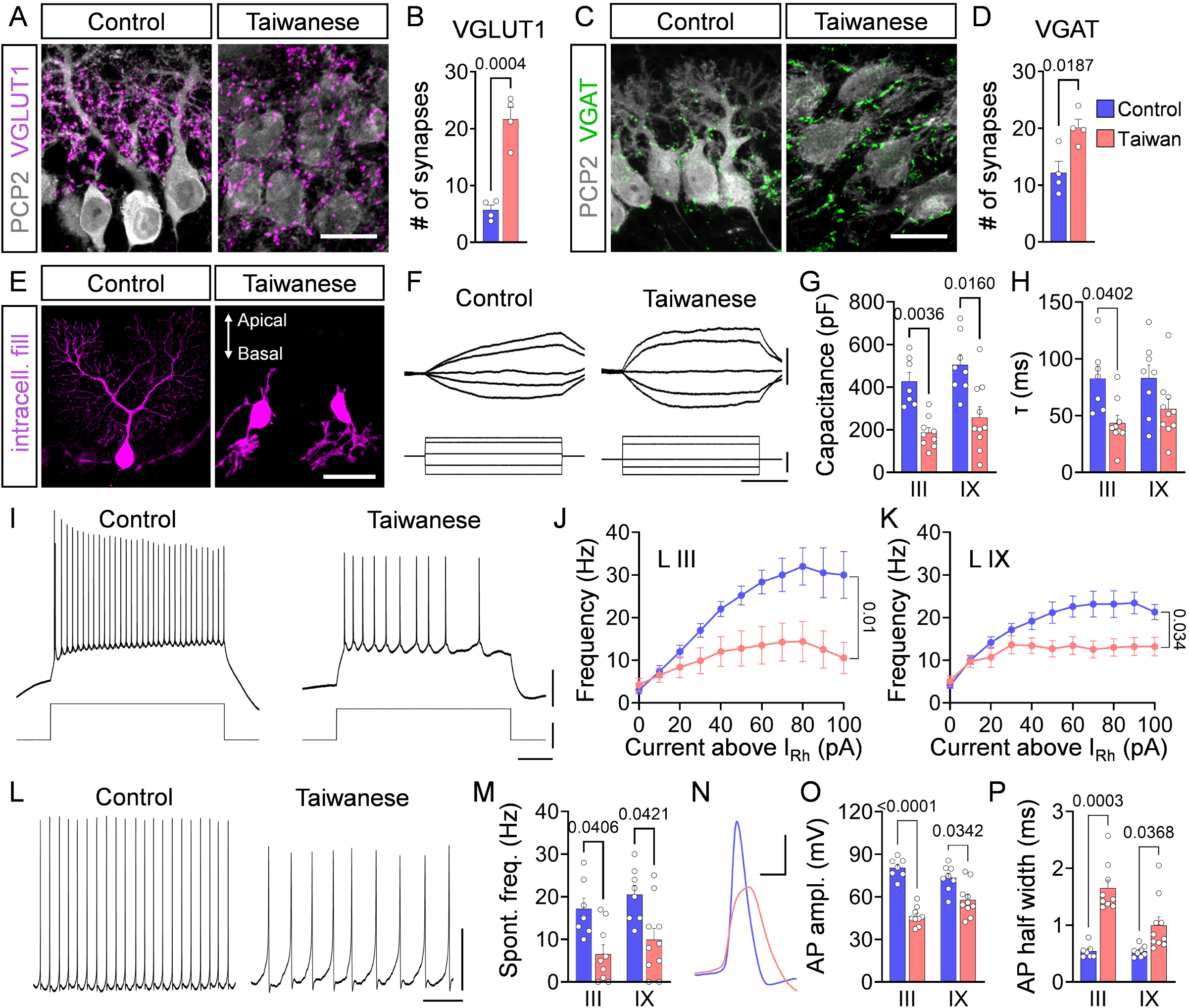
Impaired dendritic architecture, synaptic organization, and PC output in Taiwanese SMA mice. **(A)** Representative confocal images of excitatory VGLUT1⁺ synapses (magenta) onto PCP2⁺ PCs (grey). **(B)** Quantification of VGLUT1⁺ synapses onto PCs. **(C)** Representative confocal images of inhibitory VGAT⁺ synapses (green) onto PCP2⁺ PCs (grey). **(D)** Quantification of VGAT⁺ synapses onto PCs. Scale bars, 20 μm (A, C). N = 4 control and N = 4 Taiwanese SMA mice. **(E)** Confocal images of PCs filled with Atto (magenta) during whole-cell recordings from P10 control and Taiwanese SMA mice. Scale bar, 50 µm. **(F)** Representative membrane responses of PCs in lobule III following current injections. Scale bars, 5 mV, 20 pA, 100 ms. **(G, H)** Quantification of membrane capacitance (G) and membrane time constant (τ) (H) of PCs in lobules III and IX. **(I)** Representative lobule III PC firing traces during step current injections. Scale bars, 20 mV, 100 pA, 200 ms. **(J, K)** Quantification of firing frequency (frequency–current (F–I) relationship) of PCs in lobule III (J) and lobule IX (K). **(L)** Representative traces of spontaneous lobule III PC firing. Scale bars, 20 mV, 200 ms. **(M)** Quantification of spontaneous firing frequency of PCs in lobules III and IX. **(N)** Representative lobule III action potential (AP) waveform. Scale bar, 20 mV, 1 ms. **(O, P)** Quantification of AP amplitude (O) and AP half-width (P) of PCs in lobules III and IX. LIII: n = 7 control and n = 9 Taiwanese SMA PCs; LIX: n = 8 control and n = 10 Taiwanese SMA PCs from N = 4 mice per genotype for all analyses shown. Statistical analyses were performed using unpaired Student’s t-tests (B, D), Welch’s ANOVA followed by Dunnett’s T3 multiple-comparison test (G, H, M, O, P) and two-way repeated-measures ANOVA (J, K).

Next, we performed whole-cell patch-clamp recordings with simultaneous intracellular filling of individual PCs in lobules III and IX from control and Taiwanese SMA mice at P10 **(Fig. S2C)**. Intracellular fills revealed a profound defect in dendritic maturation. Rather than developing the elaborate dendritic arbor characteristic of mature P10 control PCs, Taiwanese PCs in anterior lobule III retained an immature morphology with only rudimentary dendritic trees and occasional basal dendritic orientation (**Fig. 2E**), closely resembling early postnatal PCs (**Fig. 1F**). Consistent with this reduction in dendritic complexity, membrane capacitance (reflecting membrane surface area) and membrane time constant (τ) were significantly reduced in both lobules **(Fig. 2F–H)**. In contrast, input resistance (Rin), rheobase, resting membrane potential (RMP), and action potential (AP) threshold remained unchanged **(Fig. S2D-G)**, indicating that intrinsic passive membrane properties were largely preserved. Despite this, both evoked and spontaneous firing rates were reduced by approximately 50% across lobules (**Fig. 2I–M** and **Fig. S2H, I**). Impaired firing was accompanied by decreased amplitude and broadening of the AP, particularly during the repolarization phase (**Fig. 2N–P** and **Fig. S2J**).

Overall, these findings demonstrate that profound impairment of PC dendritic maturation, altered synaptic organization, and markedly reduced PC output converge to produce widespread cerebellar circuit dysfunction in Taiwanese SMA mice.

### Distinct p53 activation patterns link developmental and degenerative cerebellar pathology across severe SMA

Previous work established that upregulation and amino-terminal phosphorylation of p53 are converging events that drive both PC and MN degeneration in the severe SMNΔ7 model.^4,10,16,37,48^ To determine whether cerebellar pathology in the Taiwanese model is similarly associated with p53 pathway activation, we examined cerebellar p53 expression between birth and end stage. In contrast to SMNΔ7 mice,^16^ p53 activation was absent from PCs throughout disease progression (**Fig. 3A, B**). Instead, robust p53 accumulation was detected throughout the external granular layer (EGL) (**Fig. 3A, B**), a transient developmental structure critical for postnatal cerebellar growth and PC maturation^49,50^. This was especially prominent at P1 (**Fig. 3B**), suggesting impaired granule cell development as a potential driver of cerebellar hypoplasia and disrupted cortical organization.

**Figure 3.**
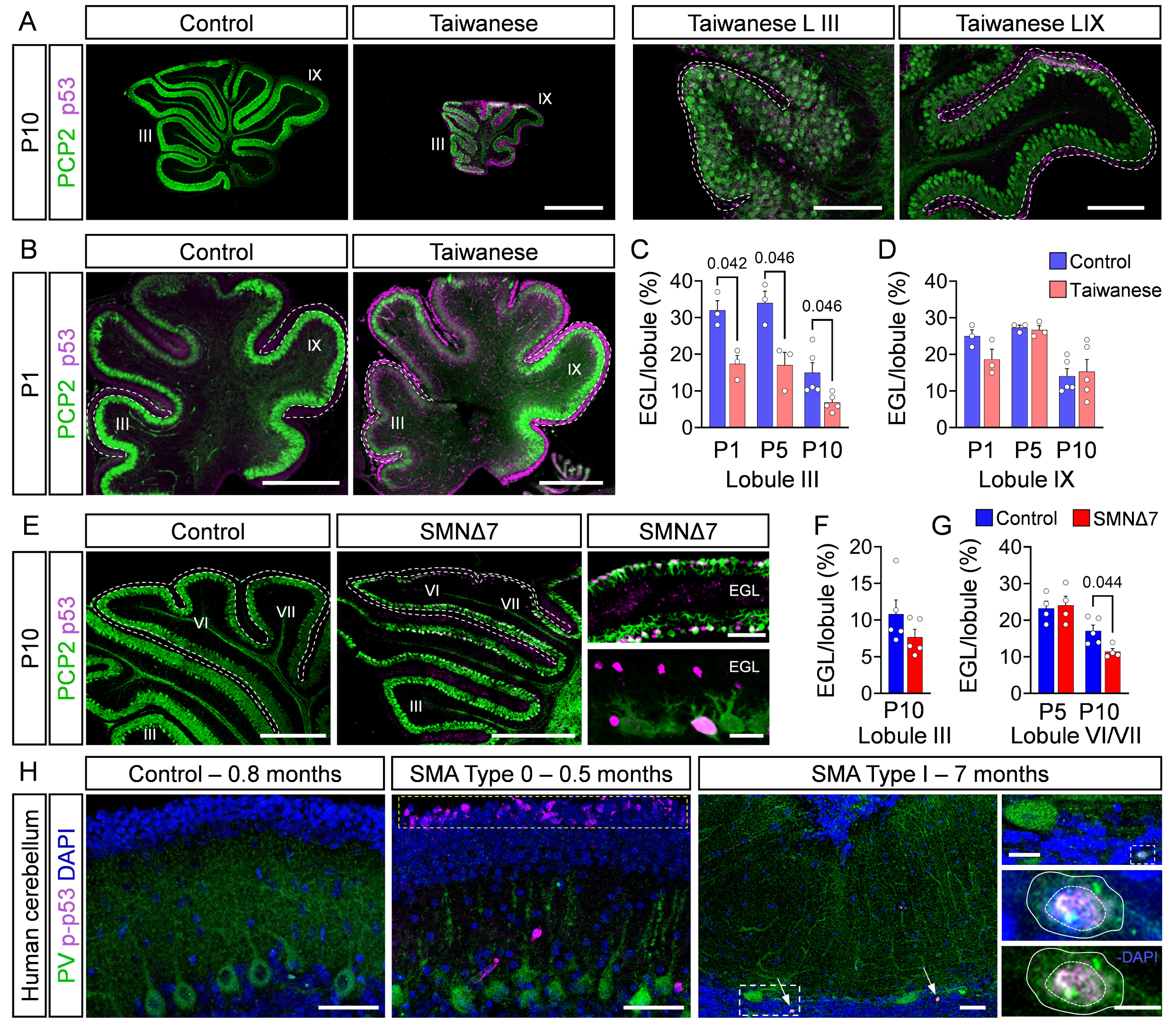
p53 activation in PCs and the EGL of severe SMA mouse models and patients. **(A)** Confocal images of sagittal cerebellar sections from P10 control and Taiwanese SMA mice immunostained for p53 (magenta) and PCP2⁺ PCs (green). Higher-magnification views of lobules III and IX are shown on the right. Scale bar, 1 mm; magnified images, 200 μm. **(B)** Representative confocal images of p53 and PCP2 immunostaining in P1 control and Taiwanese SMA mice. Scale bar, 500 μm. **(C, D)** Quantification of the EGL-to-lobule area ratio (%) in lobules III (C) and IX (D) of P1, P5, and P10 control and Taiwanese SMA mice. N = 3–4 mice per genotype and age. **(E)** Confocal images of sagittal cerebellar sections from P10 control and SMNΔ7 mice immunostained for p53 (magenta) and PCP2 (green). Higher-magnification views (right) show PCs adjacent to p53-positive EGL cells. Scale bar, 500 μm; insets, 100 μm and 20 μm. **(F, G)** Quantification of the EGL-to-lobule area ratio (%) in lobule III at P10 (F) and lobule VI/VII at P5 and P10 (G) in control and SMNΔ7 mice. N = 4–5 mice per genotype and age. **(H)** Representative confocal images of sagittal human cerebellar sections showing parvalbumin⁺ PCs (green), DAPI (blue), and p-p53^S15^ (magenta) immunostaining from a control (0.8 months), SMA Type 0 (0.5 months), and SMA Type I (7 months) patient. The yellow dashed box indicates p-p53^S15+^ EGL layer in SMA Type 0 patient. Arrows indicate p-p53^S15+^ PCs; the dashed box indicates the enlarged region; white outlines mark the PC soma and nucleus. Scale bar, 50 μm; insets, 20 μm and 5 μm. Statistical analyses were performed using multiple unpaired t-tests (C, D, G) and unpaired Student’s t-tests (F).

To determine whether EGL pathology is associated with regional developmental abnormalities, we compared the severely disorganized anterior lobule III with the more preserved posterior lobule IX. Although absolute EGL size was reduced in both lobules across developmental stages (**Fig. S3A, B**), normalization to total lobule size revealed a selective reduction in lobule III (**Fig. 3C, D**), linking EGL disruption to the pronounced developmental defects observed in vulnerable anterior cerebellar regions of the Taiwanese model.

Because SMNΔ7 mice also exhibit cerebellar hypoplasia, we investigated EGL size and re-examined p53 expression in this model. In addition to its previously reported accumulation in vulnerable PCs, p53 immunoreactivity was also detected within the EGL throughout all lobules, albeit at substantially lower levels (**Fig. 3E**). Consistent with the regional pattern of cerebellar pathology, neither absolute nor normalized EGL area was altered in the relatively resistant lobule III, whereas both measures were significantly reduced in the vulnerable lobules VI/VII (**Fig. 3E–G** and **Fig. S3C, D**). These findings indicate that EGL-associated p53 activation is a shared feature of cerebellar pathology that differs markedly in its regional distribution among different mouse models of SMA.

Amino-terminal phosphorylation of p53 is a critical activation event driving both PC and MN degeneration in the severe SMNΔ7 model.^4,10,16,37,51,52^ Because the commercial antibody used in our previous mouse studies is no longer available, murine p53^S^^18^ phosphorylation could not be assessed directly. We therefore examined the corresponding amino-terminally phosphorylated form in human SMA tissue by immunohistochemistry for p-p53^S15^. No p-p53^S15^ immunoreactivity was detected in control cerebellar tissue, whereas nuclear accumulation was observed in degenerating PCs of SMA Type I cerebellum (**Fig. 3H**), as previously reported.^16^ In striking contrast, SMA Type 0 tissue lacked p-p53^S15^ staining in PCs but exhibited prominent p-p53^S15^ immunoreactivity throughout the EGL (**Fig. 3H**). Although the available SMA Type I tissue was obtained at a substantially older age than the Type 0 sample, limiting direct comparison, p-p53^S15^ pathology was detected in both SMA tissues but was absent in age-matched control cerebella, identifying p53 phosphorylation as a conserved feature of severe human SMA.

Together, these findings support the existence of distinct profiles of p53 activation across severe forms of SMA. The PC-specific p53 upregulation observed in SMNΔ7 mice parallels the pattern detected in SMA Type I tissue, whereas EGL-associated p53 induction predominates in Taiwanese mice and is also evident in SMA Type 0 cerebellum.

### Risdiplam and AAV9-SMN result in distinct patterns of cerebellar SMN restoration

The identification of distinct cerebellar pathogenic states raised the question of whether they exhibit differential therapeutic responsiveness. We therefore investigated whether SMN-restoring therapies ameliorate cerebellar pathology in both severe SMA mouse models. Risdiplam and AAV9-SMN have previously been shown by us and other groups to robustly restore SMN in SMNΔ7 mice.^8,10,38,48,53–56^ Here, we quantified SMN protein levels in Taiwanese mice following daily intraperitoneal (IP) injection of the exon 7 splicing modifier risdiplam (3 mg/kg) or a single neonatal intracerebroventricular (ICV) injection of an adeno-associated virus serotype 9 vector expressing SMN under the GUSB promoter (AAV9-SMN; 1 × 10¹¹ genome copies). Risdiplam increased SMN protein levels from ∼10–20% to ∼50% of control in both cerebellar and spinal cord tissue (**Fig. 4A, B** and **Fig. S4A, B**). In contrast, AAV9-SMN elevated SMN to ∼120% of control in the cerebellum and ∼190% in the spinal cord of Taiwanese SMA mice (**Fig. 4A, B** and **Fig. S4A, B**). The magnitude of SMN increase achieved by both therapeutic approaches in Taiwanese mice was comparable to that we previously reported in SMNΔ7 mice.^48,53^

**Figure 4.**
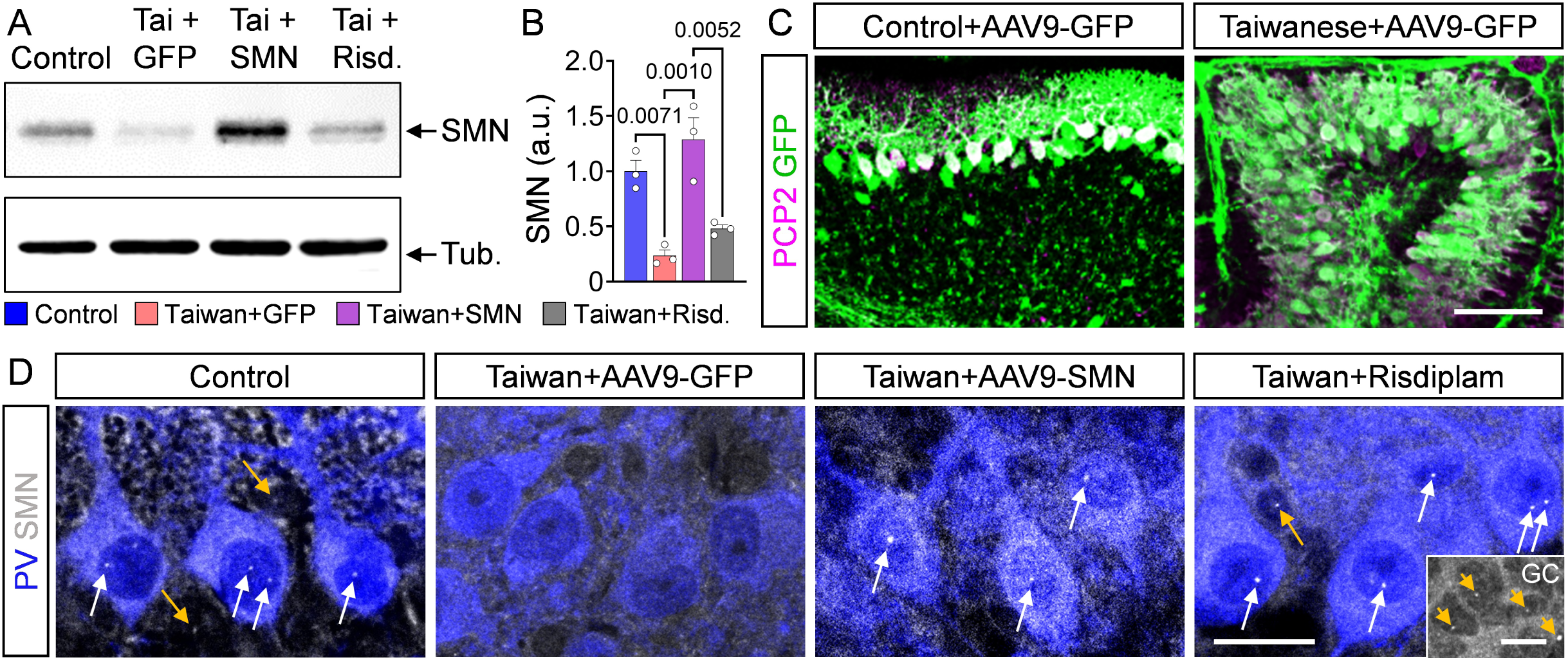
Distinct cellular patterns of cerebellar SMN restoration following risdiplam and AAV9-SMN therapy. **(A, B)** Representative Western blots (A) and quantification (B) of survival motor neuron (SMN) protein levels (tubulin as loading control) in cerebellar tissue from P10 control and Taiwanese SMA mice treated with AAV9-GFP, AAV9-SMN, or risdiplam. N = 3 mice per group. **(C)** Representative confocal images of cerebellar sections from P10 control and Taiwanese mice injected with AAV9-GFP. Sections were immunostained for PCP2 (magenta) and GFP (green). Scale bar, 100 μm. **(D)** Representative confocal images of parvalbumin (PV)^+^ PCs (blue) and SMN (grey) immunostaining in P10 control and Taiwanese SMA mice treated with AAV9-GFP, AAV9-SMN, or risdiplam. White arrows indicate SMN-containing GEMs in PCs, whereas yellow arrows indicate GEMs in non-PC cerebellar cells. Grey inset shows granule cells (GCs). Scale bar, 20 μm; GC inset, 10 μm. Statistical analyses were performed using one-way ANOVA followed by Tukey’s multiple-comparison test (B).

We next assessed vector tropism and SMN expression within the cerebellum. AAV9-GFP reporter analysis combined with GFP immunofluorescence revealed that cerebellar transduction was largely restricted to PCs, with only sparse labeling of additional cerebellar cell populations in both control and Taiwanese SMA mice (**Fig. 4C**), consistent with previous reports demonstrating preferential AAV9 tropism and an approximately 80% transduction rate.^16,57–59^ Importantly, immunohistochemistry confirmed robust SMN expression and readily detectable GEMs - nuclear structures enriched in SMN^60^ - in PCs and other cerebellar cell types throughout the cerebellar cortex of control mice, whereas GEMs were virtually absent in Taiwanese mice (**Fig. 4D**). Risdiplam treatment restored abundant GEMs throughout the cerebellar cortex, including PCs, granule cells, and cells within the molecular layer, closely resembling the distribution observed in control animals. In contrast, AAV9-SMN selectively restored GEMs in PCs, whereas GEMs remained largely absent from other cerebellar cell populations (**Fig. 4D**), consistent with the PC-centered tropism observed following AAV9-GFP administration (**Fig. 4C**).

Overall, these findings demonstrate that both therapeutics effectively restore SMN expression in Taiwanese SMA mice but exhibit distinct patterns of cellular targeting in the cerebellum.

### Risdiplam and AAV9-SMN have differential effects on cerebellar pathology in severe SMA mouse models

Having confirmed effective SMN restoration by risdiplam and AAV9-SMN in SMA mice, we next investigated their therapeutic effects on cerebellar pathology in two severe SMA mouse models. First, we investigated their therapeutic consequences in the severe SMNΔ7 model. SMNΔ7 mice were treated with either systemic IP injection of risdiplam (or the risdiplam analog SMN-C8) or neonatal ICV delivery of AAV9-SMN as previously described.^8,38^ Because SMN-C8 and risdiplam produced indistinguishable effects in this study, the corresponding data were pooled. Both AAV9-SMN and risdiplam/SMN-C8 significantly improved body weight and motor performance of SMNΔ7 SMA mice compared with their respective vehicle-treated SMA controls (AAV9-GFP and DMSO; **Fig. S5A–D**), consistent with previous reports.^10,54–56^ Because both vehicle groups showed indistinguishable phenotypes and cerebellar pathology, they were pooled for all subsequent morphological analyses and are referred to as “vehicle-treated” SMA mice. At P10–P11, both treatments restored the size of affected lobules, rescued EGL development, enhanced PC dendritic arborization, and prevented degeneration of vulnerable PCs in lobules VI/VII (**Fig. 5A, C** and **Fig. S5E–H**). This rescue was accompanied by suppression of p53 induction in both PCs and the EGL (**Fig. 5A, B**). At the circuit level, however, the extent of synaptic rescue differed between treatments. Although both treatments restored the loss of inhibitory VGAT⁺ inputs, only risdiplam therapy normalized the number of excitatory VGLUT1⁺ parallel fiber synapses, which remained significantly reduced following AAV9-SMN treatment (**Fig. 5D-G**). Together, these findings demonstrate that AAV9-SMN largely rescues intrinsic PC pathology in SMNΔ7 mice but not cerebellar circuit pathology, which requires broader cellular SMN restoration as achieved by systemic risdiplam treatment.

**Figure 5.**
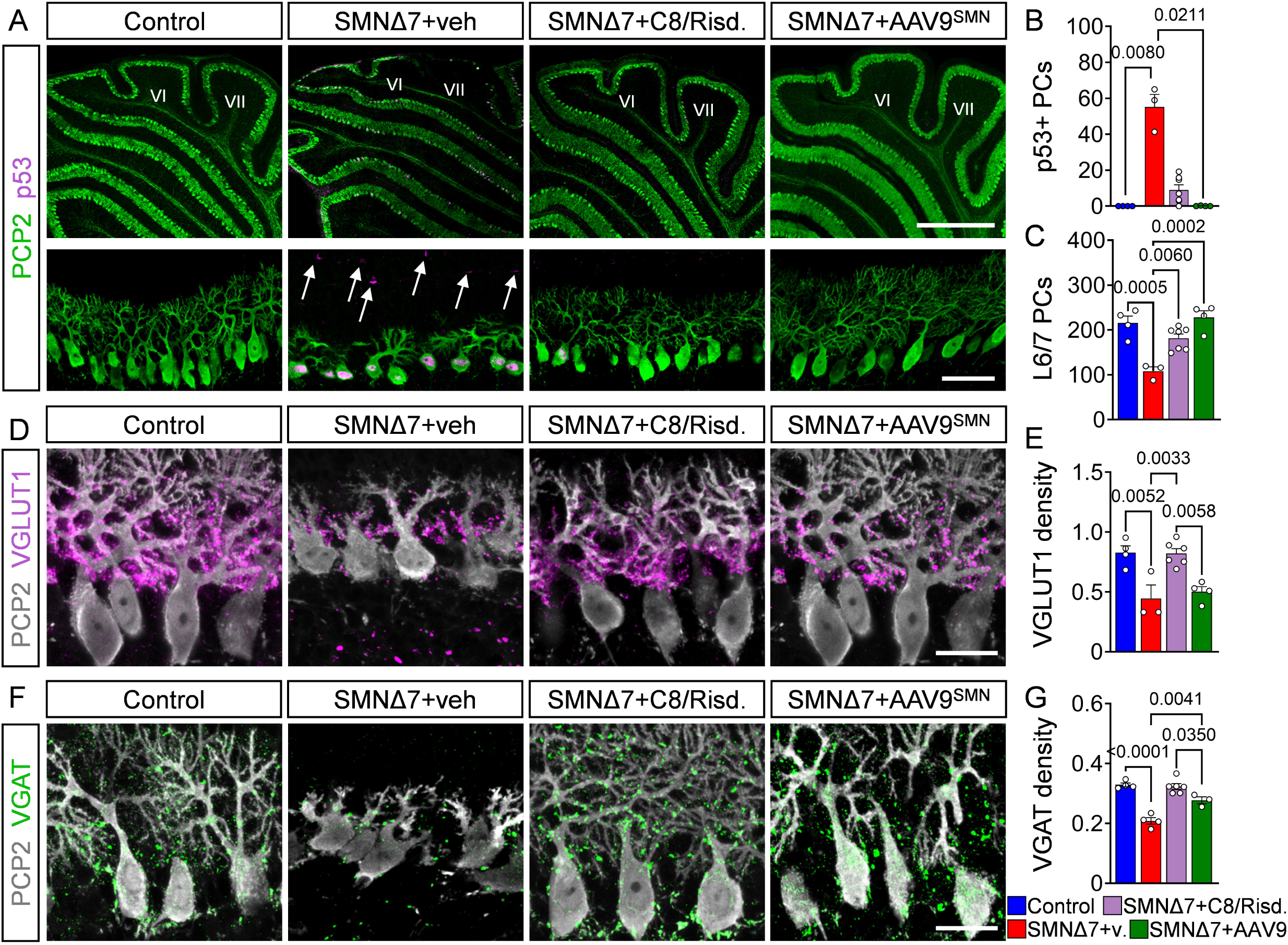
Shared and distinct effects of individual SMA therapies on cerebellar pathology. **(A)** Representative confocal images of PCP2 (green) and p53 (magenta) immunostaining in lobules VI/VII of P10 control, SMNΔ7 + vehicle (DMSO), SMNΔ7 + C8/risdiplam, and SMNΔ7 + AAV9-SMN mice. Upper panels show overview images of lobules VI/VII, whereas lower panels show higher-magnification views of PCs and the molecular layer. Arrows indicate p53-positive cells within the EGL. Scale bar, 500 μm; magnified images, 50 μm. **(B)** Quantification of p53-positive PCs (%) in P10 control (N = 4), SMNΔ7 + vehicle (N = 3), SMNΔ7 + C8/risdiplam (N = 7), and SMNΔ7 + AAV9-SMN (N = 4) mice. **(C)** Quantification of PC numbers in lobules VI/VII from P10 control (N = 4), SMNΔ7 + vehicle (N = 4), SMNΔ7 + C8/risdiplam (N = 7), and SMNΔ7 + AAV9-SMN (N = 4) mice. **(D)** Representative confocal images of PCP2 (grey) and VGLUT1 (magenta) immunostaining in lobules VI/VII from P10 control, SMNΔ7 + vehicle, SMNΔ7 + C8/risdiplam, and SMNΔ7 + AAV9-SMN mice. Scale bar, 20 μm. **(E)** Quantification of VGLUT1⁺ synaptic density on PC dendrites in P10 control (N = 4), SMNΔ7 + vehicle (N = 3), SMNΔ7 + C8/risdiplam (N = 6), and SMNΔ7 + AAV9-SMN (N = 4) mice. **(F)** Representative confocal images of PCP2 (grey) and VGAT (green) immunostaining in lobules VI/VII from P10 control, SMNΔ7 + vehicle, SMNΔ7 + C8/risdiplam, and SMNΔ7 + AAV9-SMN mice. Scale bar, 20 μm. **(G)** Quantification of VGAT⁺ synaptic density on PC dendrites in P10 control (N = 4), SMNΔ7 + vehicle (N = 4), SMNΔ7 + C8/risdiplam (N = 6), and SMNΔ7 + AAV9-SMN (N = 3) mice. Statistical analyses were performed using the Kruskal–Wallis test followed by Dunn’s multiple-comparison test (B) and one-way ANOVA followed by Tukey’s multiple-comparison test (C, E, G).

We next assessed the therapeutic effects of risdiplam and AAV9-SMN in the Taiwanese SMA model. AAV9-GFP-treated Taiwanese mice exhibited motor deficits, reduced body weight, and a lifespan of approximately two weeks (**Fig. 6A-C**), closely resembling untreated mutants^4^. Both risdiplam and AAV9-SMN treatment restored body weight and motor performance to a similar extent during early postnatal life (**Fig. 6A-C**), indicating comparable initial therapeutic efficacy. However, whereas risdiplam-treated mice continued to develop normally, the beneficial effects of AAV9-SMN were only transient. Following an initial improvement, AAV9-SMN-treated Taiwanese mice progressively failed to gain weight and developed severe motor coordination deficits characterized by near-immediate rotarod failure, postural instability, and tremor, ultimately necessitating humane euthanasia at approximately one month of age (**Fig. 6A–D, Video S1**). Importantly, the same AAV9-SMN treatment in SMNΔ7 mice does not show these later behavioral phenotypes (**Fig. 6C, D**), excluding insufficient viral potency or toxicities as potential drivers.

**Figure 6.**
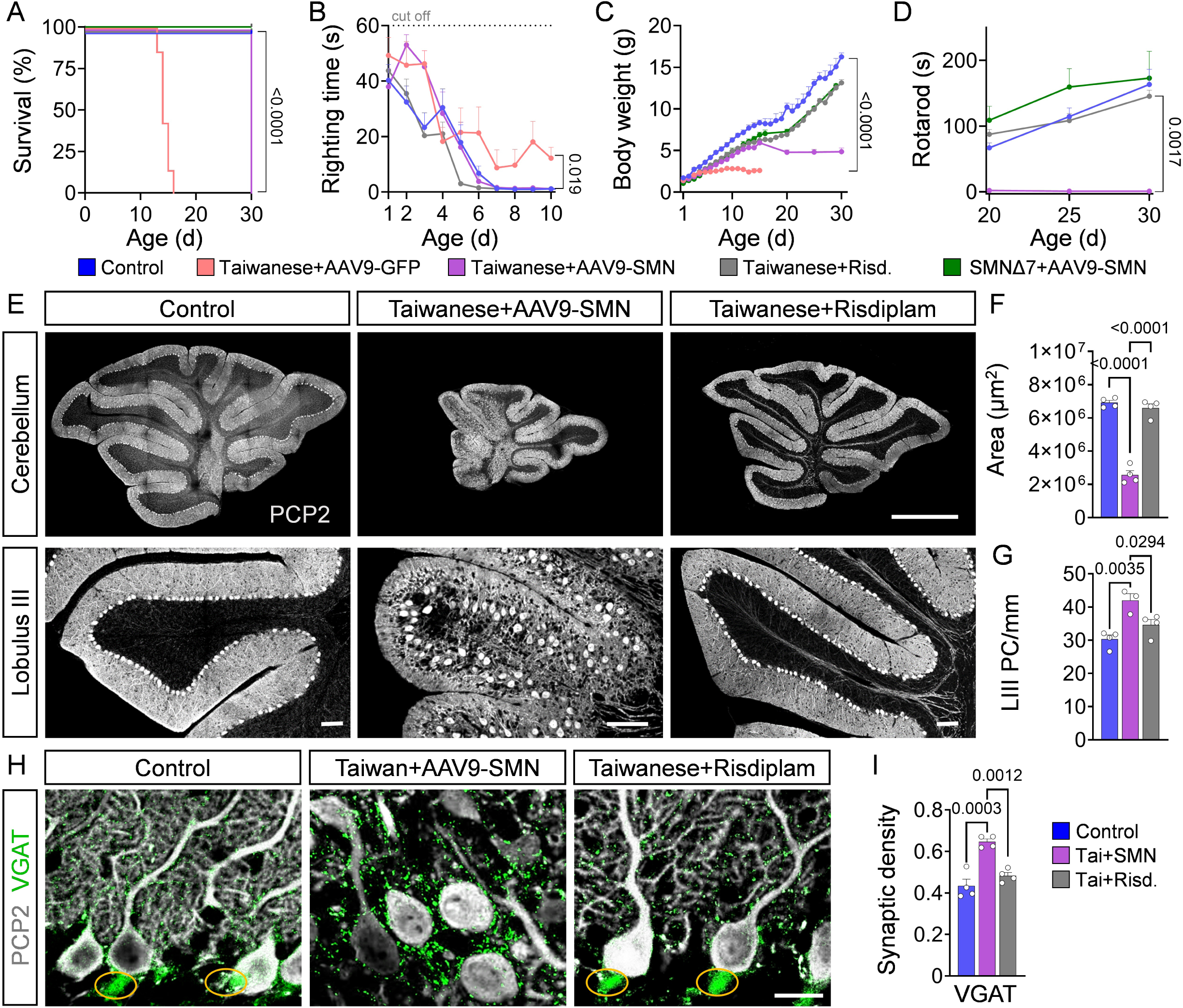
Incomplete cerebellar rescue underlies persistent ataxia-like dysfunction following AAV9-SMN therapy in Taiwanese mice. **(A–D)** Survival (A), righting time (B), body weight (C), and rotarod performance (D) from P1 to P30 in control mice, Taiwanese SMA mice treated with AAV9-GFP, AAV9-SMN, or risdiplam, and SMNΔ7 mice treated with AAV9-SMN. N = 14 control, N = 6 Taiwanese SMA + AAV9-GFP, N = 6 Taiwanese SMA + AAV9-SMN, N = 14 Taiwanese SMA + risdiplam, and N = 4 SMNΔ7 + AAV9-SMN mice. **(E)** Confocal images of sagittal cerebellar sections (top) and higher-magnification images of anterior lobule III (bottom) immunostained for PCP2 to visualize PCs in P30 control mice and Taiwanese SMA mice treated with AAV9-SMN or risdiplam. Scale bars, 1 mm (top) and 100 μm (bottom). **(F)** Quantification of total cerebellar area. **(G)** Quantification of PC density in lobule III. **(H)** Representative confocal images of inhibitory VGAT⁺ synapses (green) onto PCP2⁺ PCs (grey) in lobule III. Yellow circles indicate pinceau synapses. Scale bar, 20 μm. **(I)** Quantification of dendritic VGAT⁺ synaptic density (0–50 μm from the soma) in lobule III. For analyses shown in panels F–I, N = 4 control, N = 3 Taiwanese SMA + AAV9-SMN, and N = 4 Taiwanese SMA + risdiplam mice. Statistical analyses were performed using the log-rank Mantel–Cox test (A), mixed-effects model (REML) (B, C), two-way ANOVA followed by Tukey’s multiple-comparison test (D), and one-way ANOVA followed by Tukey’s multiple-comparison test (F, G, I).

To identify the basis of this ataxia-like behavior, we assessed central and peripheral motor circuits. Both AAV9-SMN- and risdiplam-treated Taiwanese mice showed no evidence of MN degeneration and exhibited rescue of the previously described proprioceptive deafferentation and muscle-specific NMJ denervation^4,27^ at 1 month of age (**Fig. S6A-H**), indicating effective restoration of spinal sensorimotor circuitry. In striking contrast, cerebellar pathology diverged markedly in one-month-old Taiwanese SMA mice treated with the two different SMN-restoring therapies. Whereas risdiplam rescued all major cerebellar abnormalities, AAV9-SMN-treated mice retained profound cerebellar hypoplasia, with an approximately 70% reduction in cerebellar size and preservation of the characteristic anterior-to-posterior gradient of vulnerability observed in untreated Taiwanese mice (**Fig. 6E, F**). Moreover, robust p53 expression persisted throughout the internal granular layer following AAV9-SMN treatment but was absent after risdiplam administration (**Fig. S6I**). Consistent with persistent decreased cerebellar size, PC density remained elevated despite reduced PC numbers in vulnerable anterior lobules (**Fig. 6G** and **Fig. S6J–L**). A distinct AAV9-SMN vector with the clinically relevant CBA promoter driving SMN expression similarly failed to rescue cerebellar hypoplasia (**Fig. S6N**), arguing against promoter-specific effects.

At the circuit level, AAV9-SMN-treated Taiwanese mice retained markedly reduced PC dendritic arbors, although the remaining proximal dendrites permitted synaptic quantification (**Fig. 6H**). While excitatory VGLUT1⁺ and VGLUT2^+^ inputs in AAV9-SMN-treated Taiwanese were similar to controls (**Fig. S6M**), VGAT⁺ inhibitory inputs remained abnormally increased selectively in the anterior lobule III (**Fig. 6H, I** and **Fig. S6M**), consistent with a persistent redistribution of inhibitory synapses towards the proximal dendritic compartment as a consequence of the reduced dendritic arborization. Additionally, pinceau structures formed by basket-cell terminals at the axon initial segment of PCs were absent (**Fig. 6H**). Because these specialized inhibitory synapses are critical for the temporal precision of PC firing, their loss is consistent with impaired cerebellar output and motor coordination.

Collectively, these findings demonstrate that restoration of spinal motor circuits alone is insufficient for complete functional recovery in severe SMA. Whereas viral SMN restoration effectively rescues the predominantly PC-centered cerebellar pathology of SMNΔ7 mice, broader cerebellar SMN restoration is required to prevent persistent cerebellar pathology and ataxia-like motor deficits in Taiwanese SMA mice.

## Discussion

Although SMA is classically defined as a MN disease, increasing evidence suggests that dysfunction of widespread neural circuits contributes to the disease manifestation and therapeutic outcome.^4–17^ Whether the cerebellar pathology represents a conserved feature of severe SMA and influences therapeutic recovery has remained unclear. Here, we demonstrate that cerebellar pathology is a conserved feature of severe SMA across independent mouse models and patients. However, its cellular manifestation differs markedly, with PC-intrinsic degeneration predominating in SMNΔ7 mice and SMA Type I and EGL-associated developmental pathology in Taiwanese mice and SMA Type 0. Notably, we show that distinct SMN-restoring therapies exhibit differential efficacy in rescuing cerebellar pathology, indicating that effective disease rescue depends not only on the extent but also on the proper spatial restoration of SMN. Together, these findings identify the cerebellum as a critical determinant of functional recovery in SMA mice and support a broader disease framework in which severe SMA reflects vulnerability across different motor circuits.

Previous studies reported cerebellar pathology in the severe SMNΔ7 mouse model but not in a milder SMA model,^13,16,21^ raising the question of whether cerebellar involvement is a conserved feature of severe SMA. The identification of profound cerebellar pathology in the genetically distinct Taiwanese SMA model demonstrates that cerebellar involvement is not model-specific but represents a broader feature of severe SMA. Both models exhibit cerebellar hypoplasia, PC loss and impaired cerebellar cortical output,^13,16,21^ but differ markedly in the underlying cellular pathology. SMNΔ7 mice exhibit selective PC degeneration and hypoplasia restricted to lobules VI/VII^16^, whereas the Taiwanese model displays broader hypoplasia, reduced PC numbers, and marked PC layer disorganization, prominently affecting lobules I–VII. At the circuit level, despite preserved VGLUT2⁺ climbing fiber innervation, the two models also exhibit fundamentally different synaptic phenotypes. Whereas VGLUT1⁺ excitatory and VGAT⁺ inhibitory synapses are reduced on PC dendrites in SMNΔ7 mice,^16^ Taiwanese mice display increased somatic VGLUT1⁺ and VGAT⁺ inputs. This selective pattern argues against generalized synaptic remodeling and is instead consistent with abnormal synaptic compartmentalization resulting from arrested dendritic maturation. During normal postnatal development, elaboration of the PC dendritic arbor establishes distinct excitatory and inhibitory synaptic domains,^46,47^ and impaired dendritic growth would therefore be expected to alter the spatial distribution of these inputs. Despite these differences, both mouse models show broadened APs and reduced PC firing, with impaired cerebellar cortical output.^16^ Notably, this electrophysiological signature parallels deficits observed in SMA spinal MNs,^5,8^ suggesting that distinct pathological processes converge on impaired neuronal output across multiple SMA-affected motor circuits.

Mechanistically, the different cerebellar phenotypes of the two severe SMA models are associated with distinct patterns of p53 activation. In SMNΔ7 mice, upregulation and amino-terminal phosphorylation of p53 drive cell-autonomous PC death.^16^ While selective restoration of SMN in PCs alleviates cerebellar hypoplasia in SMNΔ7 mice, p53 knockdown prevents PC death without rescuing overall cerebellar size,^16^ indicating that normal cerebellar development requires SMN-dependent functions in PCs that extend beyond survival. In addition, moderate p53 activation was also observed in the EGL of SMNΔ7 mice. By contrast, the Taiwanese model exhibits widespread p53 activation throughout the EGL while lacking detectable p53 expression in PCs, suggesting that the primary pathogenic process originates within the EGL. As the EGL serves as the transient germinal zone for cerebellar granule cell precursors and is essential for postnatal cerebellar growth and foliation,^49,50^ impaired EGL development would be expected to reduce the generation of mature granule cells within the internal granular layer. This, in turn, is likely to disrupt the reciprocal developmental interactions between granule cells and PCs required for normal PC alignment, dendritic maturation and circuit assembly, providing a plausible explanation for the profound dendritic hypoplasia and abnormal synaptic organization observed in Taiwanese mice. The same mechanism may also contribute to the reduction in PC number, as impaired granule cell support could compromise postnatal PC maturation and survival in the absence of cell-autonomous p53 activation. The coexistence of a 40% reduction in total PC number despite increased PC density further indicates that impaired cerebellar expansion is a major determinant of the phenotype. Importantly, human cerebellar pathology is broadly consistent with these observations. Whereas SMA Type I tissue exhibits p53-positive degenerating PCs resembling the SMNΔ7 phenotype,^16^ the SMA Type 0 cerebellum displays partial disorganization and increased density of PCs associated with p53 activation in the EGL, resembling the phenotype observed in Taiwanese mice. Overall, the absence of p53 activation in control cerebella supports the conclusion that both PC- and EGL-associated p53 pathology represent disease-related features of severe SMA in patients and mice. Moreover, these abnormalities were detected shortly after birth, supporting the concept that the cerebellar pathology is an early developmental feature of severe SMA.

The divergent cellular mechanisms underlying cerebellar pathology translated into distinct requirements for therapeutic rescue. In SMNΔ7 mice, AAV9-SMN suppresses p53 activation in PCs and EGL cells, rescues cerebellar growth, prevents PC degeneration, and largely restores cerebellar architecture despite incomplete normalization of excitatory synaptic organization. This closely resembles the phenotypic rescue achieved by selective SMN restoration in PCs,^16^ further supporting a PC-centered disease mechanism and indicating that neonatal AAV9-mediated SMN delivery effectively targets the critical pathogenic cell population in this model. By contrast, the same approach fails to rescue cerebellar development in Taiwanese mice, resulting in persistent dendritic hypoplasia, PC loss, and abnormal synaptic organization. This indicates that correcting SMN deficiency in PCs alone is insufficient to support normal cerebellar development but requires SMN restoration across multiple cell populations in this model. Consistent with this interpretation, systemic risdiplam completely prevents cerebellar pathology in both SMA mouse models. Overall, our findings reveal that therapeutic response is governed by the ability of individual treatments to address distinct cellular and spatial deficits that are induced by SMN deficiency during cerebellar development and are dependent on disease severity.

The distinct therapeutic requirements for cerebellar rescue are directly reflected in behavioral outcomes. Despite incomplete restoration of excitatory cerebellar circuitry in lobules VI/VII, AAV9-SMN substantially improves gross motor behavior in SMNΔ7 mice, indicating that preventing PC loss and restoring cerebellar growth is sufficient to support basic motor function. In Taiwanese mice, failure to rescue widespread pathology of cerebellar development - especially that of the anterior lobules involved in spinocerebellar sensorimotor processing and motor coordination^61,62^ - results in ataxia-like motor deficits despite rescue of spinal motor circuits. Although restoration of spinal sensorimotor circuits significantly improves motor performance in these mice^5,8^, the persistent deficits observed here indicate that optimal functional recovery additionally requires restoration of cerebellar motor circuits. In agreement, selective SMN restoration in PCs improves motor behavior in neonatal SMA mice.^16^ Together, these findings identify incomplete cerebellar rescue as a critical determinant of persistent motor dysfunction in severe SMA.

The incomplete behavioral rescue observed in SMA mice following SMN-directed therapy mirrors the persistence of motor deficits in many treated SMA patients,^2,28^ suggesting that restoration of spinal motor circuits alone is insufficient for complete functional recovery. The identification of cerebellar abnormalities in both severe SMA mouse models and patients supports the cerebellum as a clinically relevant supraspinal structure contributing to the pathology in SMA.^13,16,19,21–26^ Consistent with our findings, postmortem analyses from SMA patients treated with the AAV9-based gene therapy onasemnogene abeparvovec demonstrated cerebellar transduction largely restricted to PCs and a small subset of molecular layer cells,^63^ suggesting that incomplete cerebellar targeting may limit functional recovery. More broadly, our findings suggest that the cellular and regional distribution of SMN restoration may be as important as the absolute level achieved. In addition to the spatial distribution of SMN restoration, the developmental timing of therapeutic intervention is likely to be equally important. Impaired prenatal motor axon development and the improved motor outcomes following prenatal SMN restoration in SMA patients support the concept that critical disease processes are already established during embryonic development.^53,64^ Consistent with this, whereas cerebellar growth and circuit assembly occur predominantly after birth in mice, comparable developmental processes begin during late gestation in humans and continue throughout infancy,^65^ raising the possibility that the therapeutic window for preventing cerebellar pathology extends into prenatal development in SMA patients. Although no approved SMN-directed therapy has yet demonstrated clear clinical superiority in the absence of direct head-to-head comparisons,^66^ future clinical studies should determine whether biomarkers of cerebellar dysfunction correlate with therapeutic efficacy and long-term motor outcomes in SMA patients. Together, these findings support a broader disease framework in which severe SMA reflects dysfunction of distributed motor networks and identify supraspinal circuit restoration as a critical determinant of durable therapeutic recovery.

## Data availability

The authors confirm that the data supporting the findings of this study are available within the article and its Supplementary material. Some data are not publicly available due to patient-related restrictions, as they contain information that could compromise the privacy of research participants. Additionally, certain derived data from mouse experiments are available from the corresponding authors upon reasonable request.

## Acknowledgements

We thank Drs. Johannes Hirrlinger, Jens Eilers, and Tobias Langenhan for their contributions of reagents and access to facilities. Some of the human tissue was obtained from the NIH Neurobiobank at the University of Maryland, Baltimore, MD, USA.

## Supplementary material

Supplementary material is available online.

