## Supplemental Figures for "Incomplete cerebellar circuit restoration limits functional recovery following SMN therapy in severe spinal muscular atrophy"

Figure S1 Structural abnormalities with preserved motor cortex morphology in the Taiwanese model

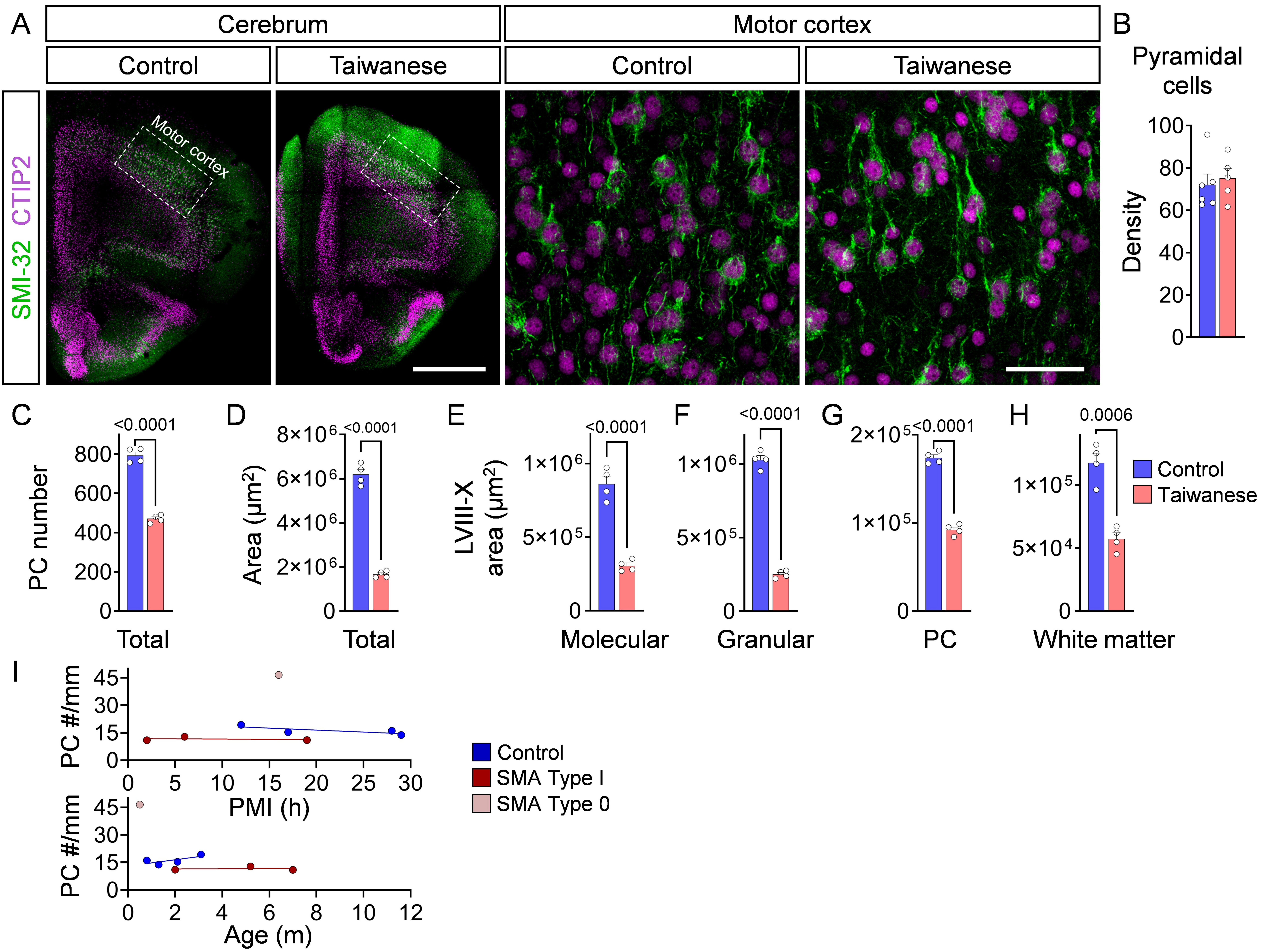

Figure S2 Membrane excitability is unaltered in PCs of the Taiwanese mouse model

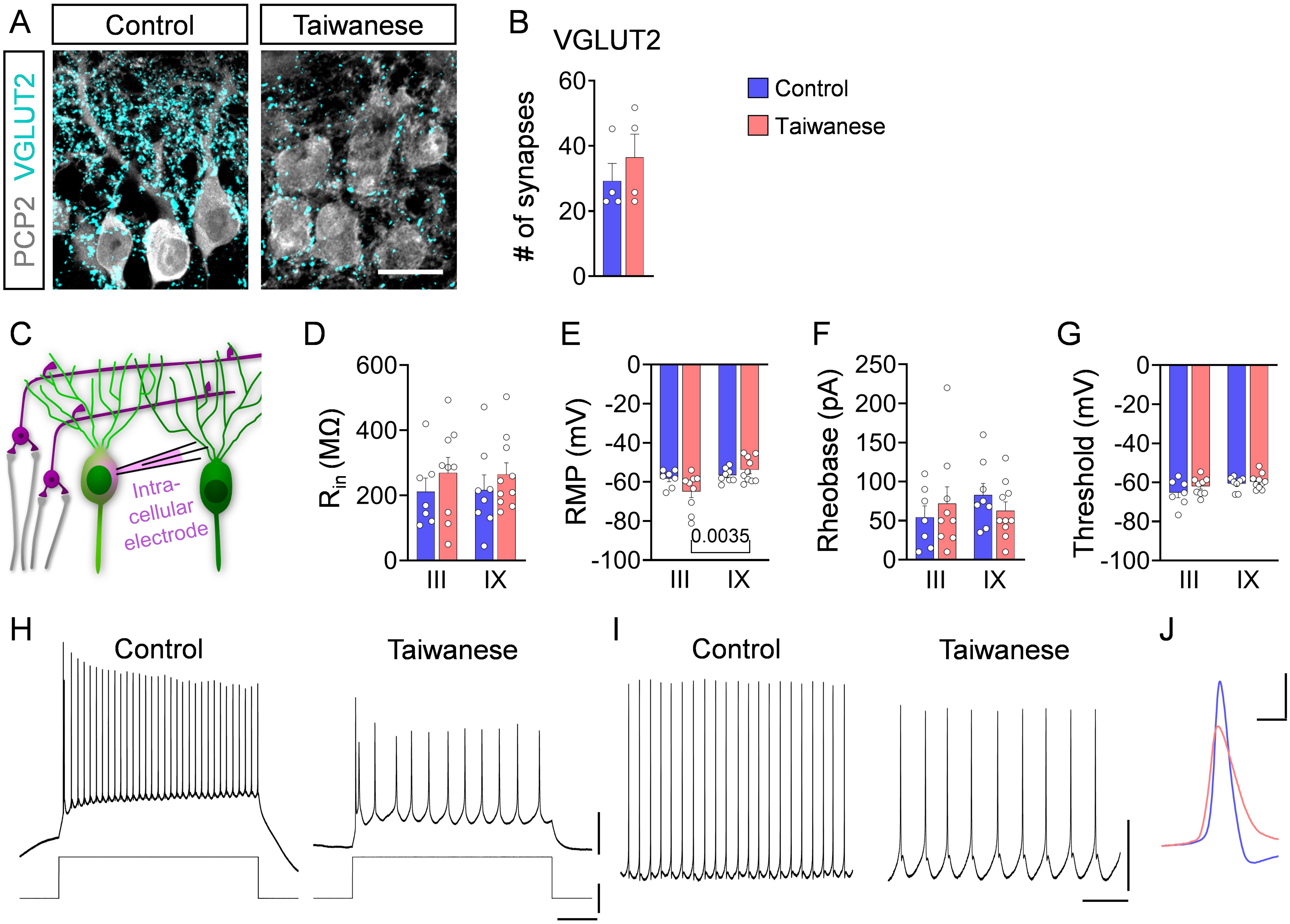

Figure S3 EGL area is severely affected in vulnerable lobules across severe SMA mouse models

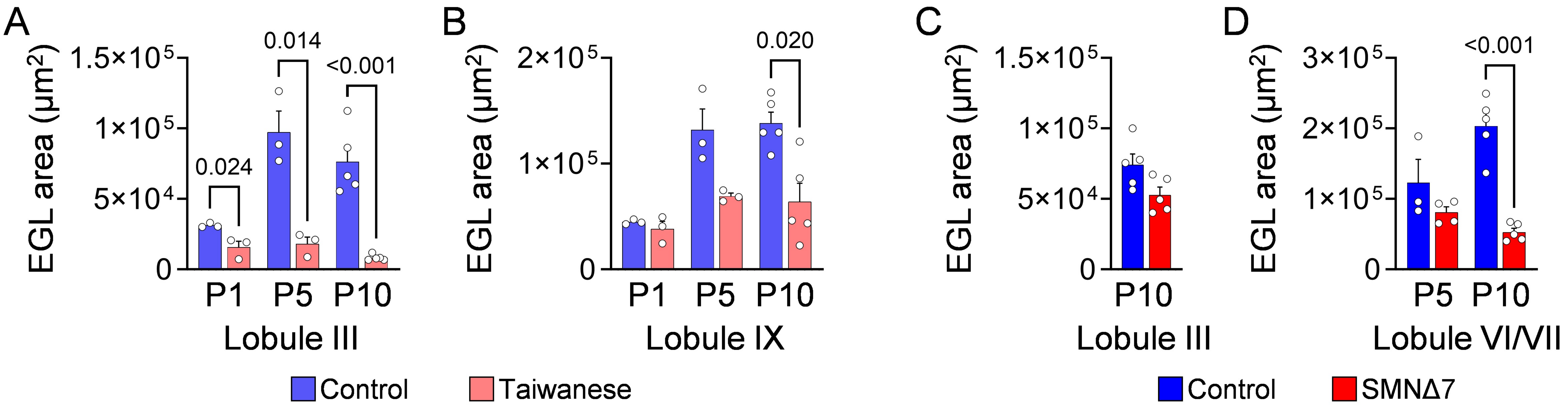

Figure S4 SMN restoration in spinal cord tissue of Taiwanese mice following risdiplam and AAV9-SMN therapy

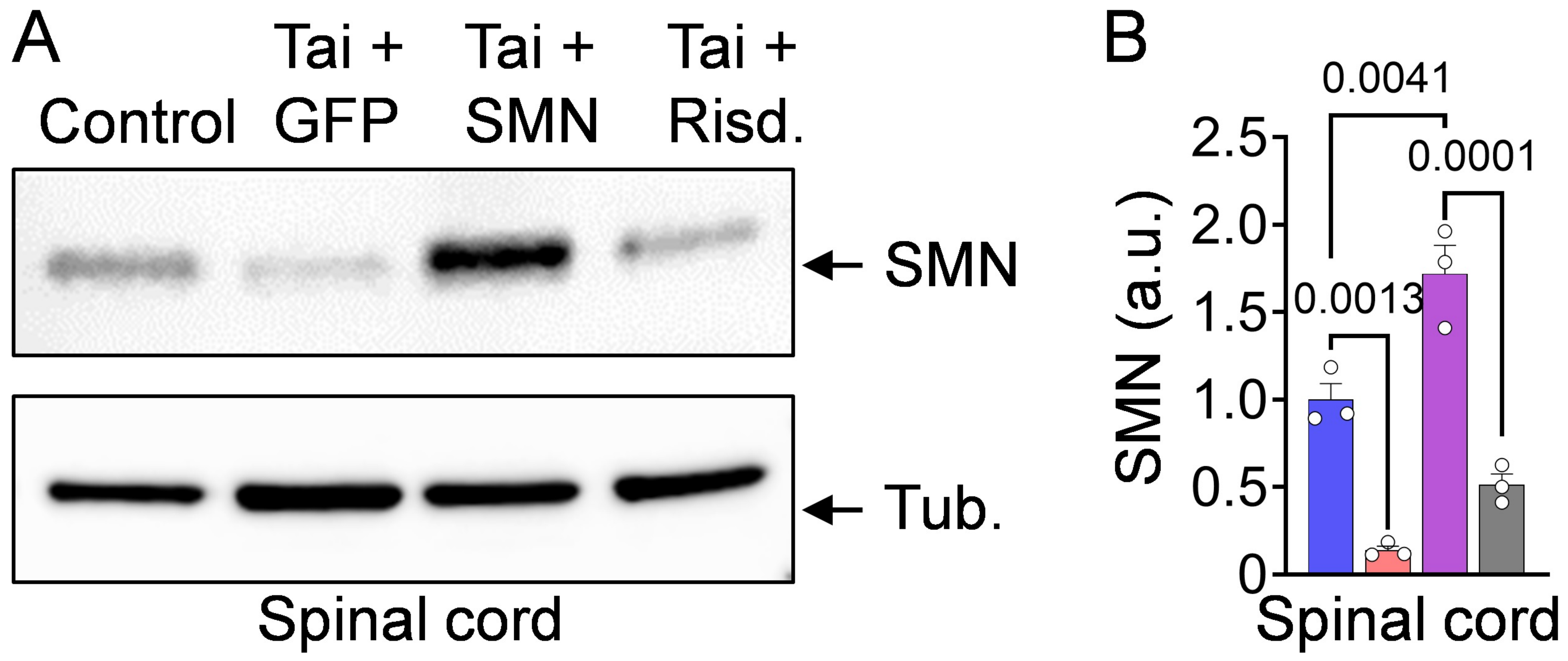

Figure S5 SMN-restoring therapies ameliorate disease phenotypes in SMNΔ7 mice

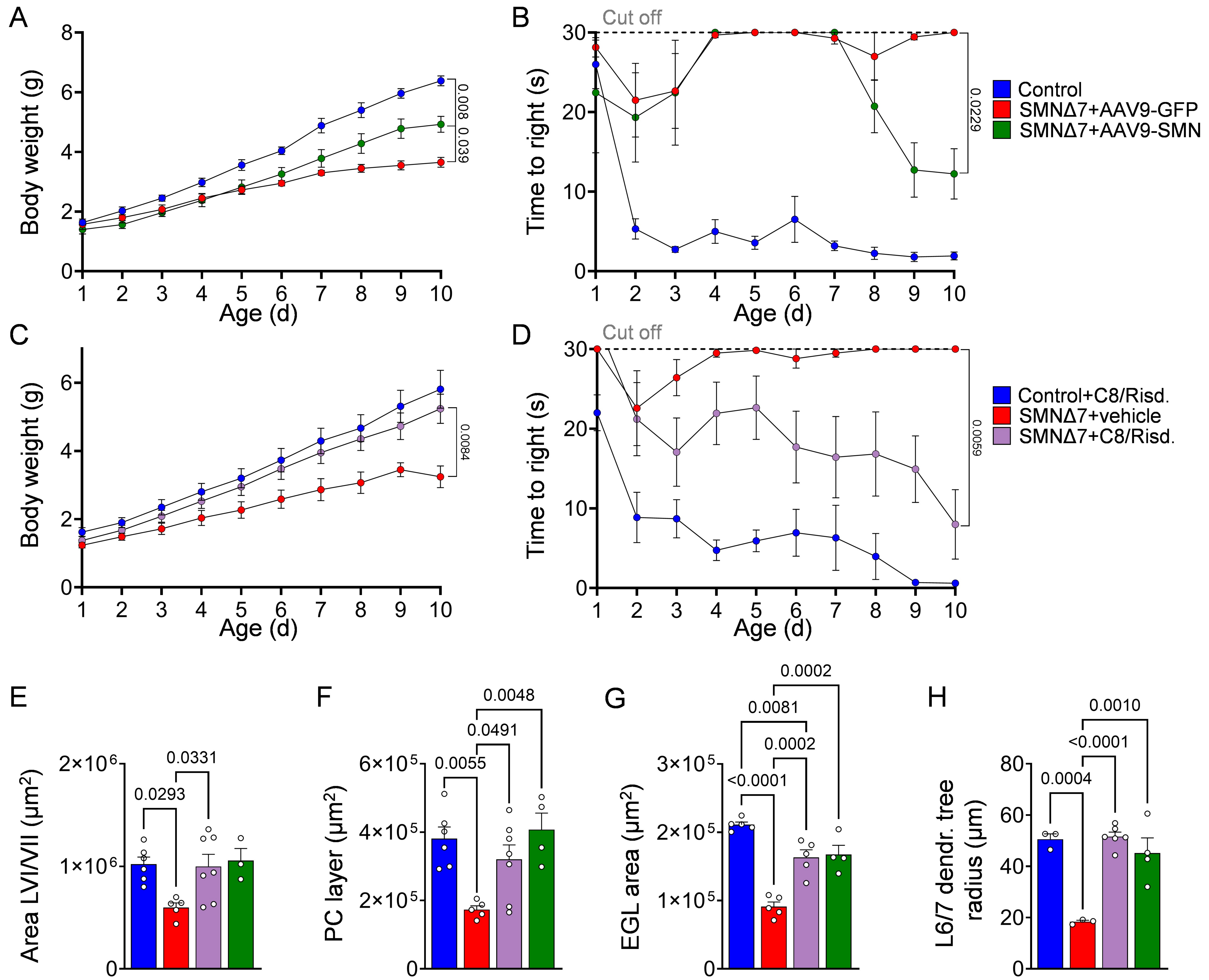

Figure S6 Both SMN therapies rescue therapy rescue sensory-motor circuit in Taiwanese model

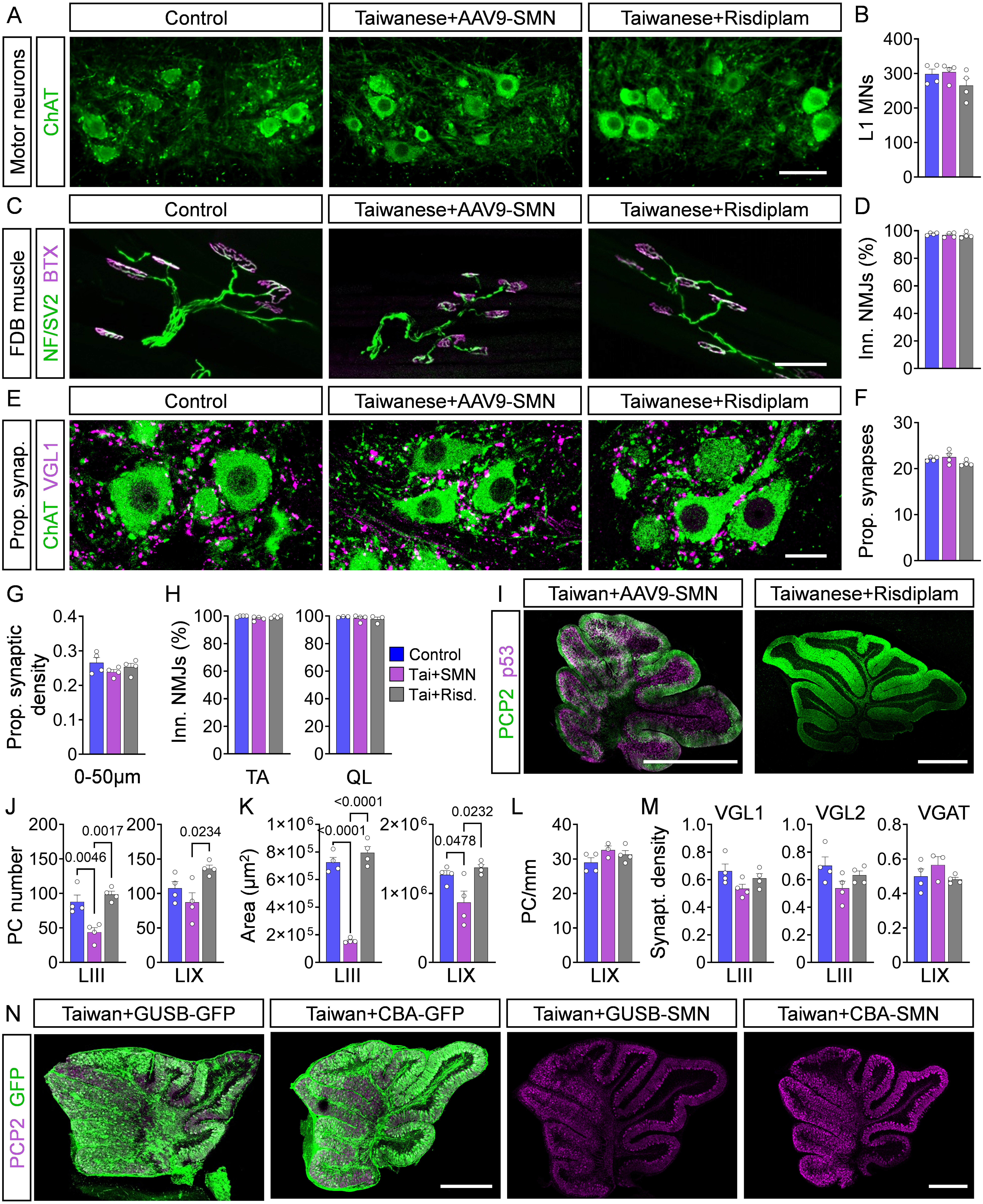
