## Supplemental Tables for "Incomplete cerebellar circuit restoration limits functional recovery following SMN therapy in severe spinal muscular atrophy"

**Supplementary Tables**

**Supplementary Table 1: List of patients.**

| **Patient ID** | **Group** | **Cause of Death** | **Sex** | **Age (m)** | **PMI (h)** | **Origin** |
| --- | --- | --- | --- | --- | --- | --- |
| Control #1 | Control | SUDI | male | 2.1 | 17 | NIH |
| Control #2 | Control | SUDI | male | 1.3 | 29 | NIH |
| Control #3 | Control | Cardiomyopathy | male | 3.1 | 12 | NIH |
| Control #4 | Control | SUDI | male | 0.8 | 28 | NIH |
| SMA #1 | SMA Type I | SMA complications | female | 2 | 19 | NIH |
| SMA #2 | SMA Type I | Respiratory failure | male | 5.2 | 6 | NIH |
| SMA #3* | SMA Type I | Refractory hypoxemia | male | 7 | 2 | JHU |
| SMA #4 | SMA Type 0 | Respiratory failure | male | 0.5 | 16 | JHU |

SUDI = sudden unexpected death in infancy, NIH = National Institute of Health, JHU = Johns Hopkins University. *Patient SMA #3 received risdiplam from 5 to 7 months of age.

**Supplementary Table 2: List of antibodies.**

| **Name** | **Company** | **Cat #** | **Host** | **Reactivity** | **Dilution** |
| --- | --- | --- | --- | --- | --- |
| PCP-2 | Santa Cruz | Sc-137064 | Mouse | Mouse | 1:400 |
| Parvalbumin | Synaptic Systems | 195 006 | Chicken | Human/mouse | 1:5,000 |
| p53 | Leica Novocastra | NCL-p53-CM5p | Rabbit | Mouse | 1:1,000 |
| p-p53^S15^ | Cell Signaling | E9Y4U | Rabbit | Human | 1:500 |
| VGLUT1 | Synaptic Systems | 135 304 | Guinea pig | Mouse | 1:5,000 |
| VGLUT2 | Synaptic Systems | 135 403 | Rabbit | Mouse | 1:1,000 |
| VGAT | Synaptic Systems | 131 004 | Guinea pig | Mouse | 1:500 |
| ChAT | Millipore | AB144P | Goat | Mouse | 1:500 |
| SMN | BD Transd. Labs | 610646 | Mouse | Mouse | 1:100 |
| SV2 | DSHB | SV2-c | Mouse | Mouse | 1:500 |
| NF-H | DSHB | 2H3-c | Mouse | Mouse | 1:1,000 |
| NF-H [SMI-32] | Biolegend | 801712 | Mouse | Mouse | 1:500 |
| NF-M | Sigma-Aldrich | AB1987 | Rabbit | Mouse | 1:1000 |
| GFP | Abcam | AB13970 | Chicken | Mouse | 1:2,000 |
| CTIP2 | Abcam | AB18465 | Rat | Mouse | 1:500 |
| Bungarotoxin | Invitrogen | B35451 | N/A | Mouse | 1:1,000 |
| DAPI | Thermo Fisher | 62248 | N/A | Human | 1:3000 |
